# Genes associated with neuropsychiatric disease increase vulnerability to abnormal deep grey matter development

**DOI:** 10.1101/342394

**Authors:** Harriet Cullen, Michelle L Krishnan, Saskia Selzam, Gareth Ball, Alessia Visconti, Alka Saxena, Serena J Counsell, Jo Hajnal, Gerome Breen, Robert Plomin, A David Edwards

**Affiliations:** Centre for the Developing Brain, Kings College, London, SE1 7EH, United Kingdom; Institute of Psychiatry, Psychology and Neuroscience, Kings College London, SE5 8AF, United Kingdom; Translational Medicine, Neuroscience and Rare Diseases, Roche Pharmaceutical Research and Early Development, Roche Innovation Center, 4070 Basel, F. Hoffmann-La Roche, Ltd.; Developmental Imaging, Murdoch Children’s Research Institute, Melbourne, Australia; Department of Twin Research and Genetic Epidemiology, King’s College London, SE1 7EH United Kingdom; NIHR Biomedical Research Centre, Guy’s and St Thomas’ NHS Foundation Trust, London SE1 9RT, United Kingdom

**Keywords:** Brain Development, Preterm, Deep Grey Matter, Polygenic Risk Score, Magnetic Resonance Imaging, Psychiatric Pathology

## Abstract

1.

**Background:** Neuropsychiatric disease has polygenic determinants but is often precipitated by environmental pressures, including adverse perinatal events. However, the way in which genetic vulnerability and early-life adversity interact remains obscure. Preterm birth is associated with abnormal brain development and psychiatric disease. We hypothesised that the extreme environmental stress of premature extra-uterine life could contribute to neuroanatomic abnormality in genetically vulnerable individuals.

**Methods:** We combined Magnetic Resonance Imaging (MRI) and genome-wide single nucleotide polymorphism (SNP) data from 194 infants, born before 33 weeks of gestation, to test the prediction that: the characteristic deep grey matter abnormalities seen in preterm infants are associated with polygenic risk for psychiatric illness. Summary statistics from a meta-analysis of SNP data for five psychiatric disorders were used to compute individual polygenic risk scores (PRS). The variance explained by the PRS in the relative volumes of four deep grey matter structures (caudate nucleus, thalamus, subthalamic nucleus and lentiform nucleus) was estimated using linear regression both for the full, mixed-ancestral, cohort and a subsample of European infants.

**Results:** The PRS was negatively associated with: lentiform volume in the full cohort (β=−0.24, p=8×10^−4^) and the European subsample (β=−0.24, p=8×10^−3^); and with subthalamic nuclear volume in the full cohort (β=−0.18, p=0.01) and the European subsample (β=−0.26, p=3×10^−3^).

**Conclusions:** Genetic variants associated with neuropsychiatric disease increase vulnerability to abnormal deep grey matter development and are associated with neuroanatomic changes in the perinatal period. This suggests a mechanism by which perinatal adversity leads to later neuropsychiatric disease in genetically predisposed individuals.

## 2. Introduction

Preterm birth accounts for around 11% of all births and is a leading cause of infant mortality and morbidity (1). It is a profound early-life stressor that is strongly associated with cognitive impairment, cerebral palsy, autism spectrum disorders and psychiatric disease (2)(3)(4)(5). Preterm birth is associated with both cerebral grey and white matter abnormalities. Deep grey matter appears to be especially vulnerable (6)(7)(6)(8) and reduced deep grey matter volumes have been associated with poorer neurodevelopmental outcome and neurodevelopmental disability (9)(10).

Neuroanatomical and functional outcomes vary significantly between individuals and are likely to be modulated by the interaction of genetic and environmental factors (11)(12). Adverse psychiatric outcomes have moderate to high heritability (13), as do many brain-imaging phenotypes (14) (15), even in early infancy (16). Genome-wide association studies (GWAS) have shown that this heritability is largely due to many genetic variants of small effect (17). GWAS have become increasingly informative as sample sizes have grown, allowing detection of small effects of single nucleotide polymorphisms (SNPs), although SNPs reaching the stringent statistical criteria for genome-wide significance (typically P < 5×10^−8^) are few and collectively explain only a small percentage of the genetic variance of the disorder in question (18). However, it is now possible to generate individual-specific genotypic scores to predict phenotypic variance. GWAS results can be used to construct a polygenic risk score (PRS), which is an aggregate of trait-related effect sizes of SNPs across the genome in independent samples (19).

We reasoned that if genes associated with psychiatric disease increase vulnerability to environmental effects on brain development, individuals with a high polygenic risk for psychiatric disease who are subjected to the extreme environmental stress of preterm birth would be more likely to develop adverse consequences. Here, we exploit the power of previous large genome-wide studies measuring the polygenic risk for five different psychiatric disorders (20), combining this with a large set of Magnetic Resonance Imaging (MRI) data in preterm infants to test the prediction that: the characteristic deep grey matter abnormalities seen in preterm infants are associated with a higher polygenic risk for psychiatric illness.

## 3. Materials and Methods

### 3.1 Subjects

Preterm infants were recruited as part of the EPRIME (Evaluation of Magnetic Resonance (MR) Imaging to Predict Neurodevelopmental Impairment in Preterm Infants) study and were imaged at term equivalent age. The EPRIME study was reviewed and approved by the National Research Ethics Service, and all infants were studied following written consent from their parents.

### 3.2 **Imaging**

#### 3.2.1 MRI acquisition

MRI was performed on a Philips 3-Tesla system (Philips Medical Systems, Best, The Netherlands) within the Neonatal Intensive Care Unit using an 8-channel phased array head coil.

T1-weighted MRI was acquired using: repetition time (TR): 17 ms; echo time (TE): 4.6 ms; flip angle 13°; slice thickness: 0.8 mm; field of view: 210 mm; matrix: 256×256 (voxel size: 0.82×0.82×0.8 mm). T2-weighted fast-spin echo MRI was acquired using: TR: 14730 ms; TE: 160 ms; flip angle 90°; field-of-view: 220 mm; matrix: 256×256 (voxel size: 0.86×0.86×2 mm) with 1mm overlap.

Pulse oximetry, temperature and heart rate were monitored throughout and ear protection was used for each infant (President Putty, Coltene Whaledent, Mahwah, NJ; MiniMuffs, Natus Medical Inc, San Carlos, CA).

#### 3.2.2 Image processing

T1-weighted images were brain-extracted with co-registered T2 brain masks (FSL’s BET; FSL 5.0.8, http://fsl.fmrib.ox.ac.uk/fsl) and corrected for bias field inhomogeneities (30). Each subjects’ T1-weighted image was aligned to a 40-week neonatal template (31) using nonlinear registration (32). Voxelwise maps of volume change induced by the transformation were characterized by the determinant of the Jacobian operator, referred to here as the Jacobian map. These maps include a global scaling factor, so Jacobian values reflect tissue volume differences due to both local deformation and global head size. T1-derived Jacobian maps were iteratively smoothed to a FWHM of 8mm (AFNI’s 3dBlurToFWHM; http://afni.nimh.nih.gov/afni). Each map was then log-transformed so that values greater than zero represent local areal expansion in the subject relative to the target and values less than zero represent areal contraction.

#### 3.2.3. Brain Endophenotype

The Jacobian maps were used to estimate the volume of regions of interest in the deep grey matter. Volume estimates for the bilateral volumes of the thalamus, subthalamic nucleus, caudate nucleus and lentiform nucleus (putamen and globus pallidus) were extracted by computing the mean log(Jacobian) within each region-specific mask. Masks were defined using a neonatal atlas (33) aligned to a 40-week template.

#### 3.3 **Genotyping and quality control**

348 saliva samples were collected using Oragene DNA OG-250 kits (DNAGenotek Inc., Kanata, Canada) and genotyped on Illumina HumanOmniExpress-24 v1.1 arrays (Illumina, San Diego, CA, USA). Individuals with genotyping completeness < 95% were excluded (29 individuals removed). SNPs with a minor allele frequency < 0.01 (24546 SNPs), a missing genotype rate < 99% (14672 SNPs) and deviations from Hardy–Weinberg equilibrium, P < 1 × 10^−7^ (2307 SNPs) were removed. Where pairs with high levels of relatedness existed (pi-hat > 0.3) only one member of each pair was retained (44 individuals removed). This resulted in a sample of 275 individuals with high-quality genetic data (635266 SNPs). All quality control steps were carried out using PLINK 1.9 (34) (Software: https://www.coggenomics.org/plink/1.9/).

### 3.4 Population Stratification and Sample Selection

After pruning to remove markers in high linkage disequilibrium (r^2^ > 0.1, 72900 SNPs retained) we performed a Principal Component Analysis using PLINK 1.9 (35). Inspection of the first two principal components in combination with reported ethnicity was used to define three ancestral populations: European, Asian and African (**Supplementary Information, Fig. S1**). Outliers from these three ancestral populations were excluded (35 individual removed). These three populations formed the cohorts for ongoing analysis (240 individuals).

Only those individual with MRI T1 data were retained (214 of the 240); individuals with marked cerebral pathology were additionally excluded (20 individuals). Our final sample comprised of 194 unrelated preterm infants (104 males, 90 females), mean gestational age (GA) 29.7 weeks, mean postmenstrual age at scan 42.6 weeks, including 122 individuals in the European cohort, 48 in the Asian cohort and 24 in the African cohort (**Table 1**).

**Table 1.** Summary statistics for three different ancestral cohorts.

| **Ancestry** | **Number**<br>**of**<br>**Subjects** | **Mean**<br>**Gestational**<br>**Age (weeks)** | **Mean**<br>**Postmenstrual**<br>**Age at Scan**<br>**(weeks)** | **Mean**<br>**Intracranial**<br>**Volume**<br>**(ml)** |
| --- | --- | --- | --- | --- |
| **European** | 122 | 29.7±0.2 | 42.3±0.2 | 569.1±5.8 |
| **Asian** | 48 | 29.6±0.3 | 42.7±0.3 | 557.1±11.0 |
| **African** | 24 | 29.3±0.6 | 43.4±0.6 | 585.9±16.0 |
| **Combined Sample** | 194 | 29.7±0.2 | 42.6±0.2 | 568.2±4.9 |

### 3.5 Polygenic Scoring

We computed polygenic risk scores (PRS) for the 194 unrelated individuals using odds ratios and *P*-values from summary statistics obtained by the primary GWAS analysis from the Cross-Disorder Group of the Psychiatric Genomics Consortium (20). This analysis combined the effects of five psychiatric diseases (autism spectrum disorder, attention deficit-hyperactivity disorder, bipolar disorder, major depressive disorder, and schizophrenia) in 33332 cases and 27888 controls of European ancestry. The authors used a meta-analytic approach that applied a weighted Z-score with weights indicating the sample-size of the disease specific studies. The greatest power was therefore for SNPs that have an effect in multiple disorders.

PRS were generated in PRSice (36). Initially the three ancestral sub-samples were considered independently. Quality-controlled SNPs were pruned for linkage disequilibrium based on *P*-value informed clumping using a r^2^ = 0.1 cut-off within a 250-kb window to create a SNP-set in linkage equilibrium for each of our ancestral cohorts. PRS were computed at five different *P*-value thresholds (PT) in the discovery GWAS summary statistics: 0.001, 0.01, 0.05, 0.1, 0.5. Scores were computed using genotyped SNPs in our target dataset. *P*-value thresholds and the number of SNPs contributing to the PRS for each of the three ancestral subsamples at each threshold are summarized in **Supplementary Table S1** in **Supplementary Information.**

To control for population stratification, we regressed the PRS on the first 10 principal components of our ancestry matrix and used the residuals in all subsequent analysis. Details of the effect of this regression on the PRS distributions for our three ancestral cohorts are outlined in (**Supplementary Information: Supplementary Methods and Supplementary Figures S2 and S3**)

### 3.6 Statistical Analysis

Bilateral deep grey matter brain volumes for the thalamus, subthalamic nucleus, caudate nucleus and lentiform nucleus (globus pallidus and putamen) were corrected for intracranial volume and gestational age at birth. Corrected volumes were then regressed on the ancestry-corrected PRS generated at five different *P*-value thresholds using simple linear regression in R. The variance explained by the PRS was obtained by squaring the regression coefficient.

Our full sample includes infants from three different ancestral cohorts: European, Asian and African. In contrast, the GWAS meta-analysis results (20) used to compute the PRS were compiled using subjects of European ancestry only. We have therefore conducted an additional sensitivity analysis using a sub-sample of our cohort comprising only the European infants.

As an additional check, we sought to confirm that the psychiatric PRS was not associated with degree of prematurity and we regressed the ancestry-corrected PRS against gestational age alone. We also looked for a possible relationship between our psychiatric PRS and developmental outcome as assessed using the Bayley Scale of Infant and Toddler Development (37).

Results quoted in **Tables 2** and **3** are raw P-values. We have additionally computed a multiple-comparison *P*-value threshold. Since both the deep grey matter volumes and the five polygenic risk score thresholds are highly correlated we have used the method proposed by (38) to compute the effective number of independent tests performed (M_e_ff). This accounts for the correlation structure between measures and calculates the M_e_ff based on the observed eigenvalue variance using the matSpD interface (https://gump.qimr.edu.au/general/daleN/matSpD/). This calculation was performed for both the deep grey matter volumes (M_eff_dgm_) and the differently-thresholded polygenic scores (M_eff_prs_). The *P*-value for significance was determined as 0.05 divided by the product of M_eff_dgm_ and M_eff_prs_. The four deep grey matter volumes resulted in two independent tests and the five differently-thresholded polygenic scores three independent tests, giving a multiple-comparison corrected *P*-value threshold of p < 0.008333. Results that survive the multiple-comparison correction in **Tables 2** and **3** are indicated with a **\***.

**Table 2.**
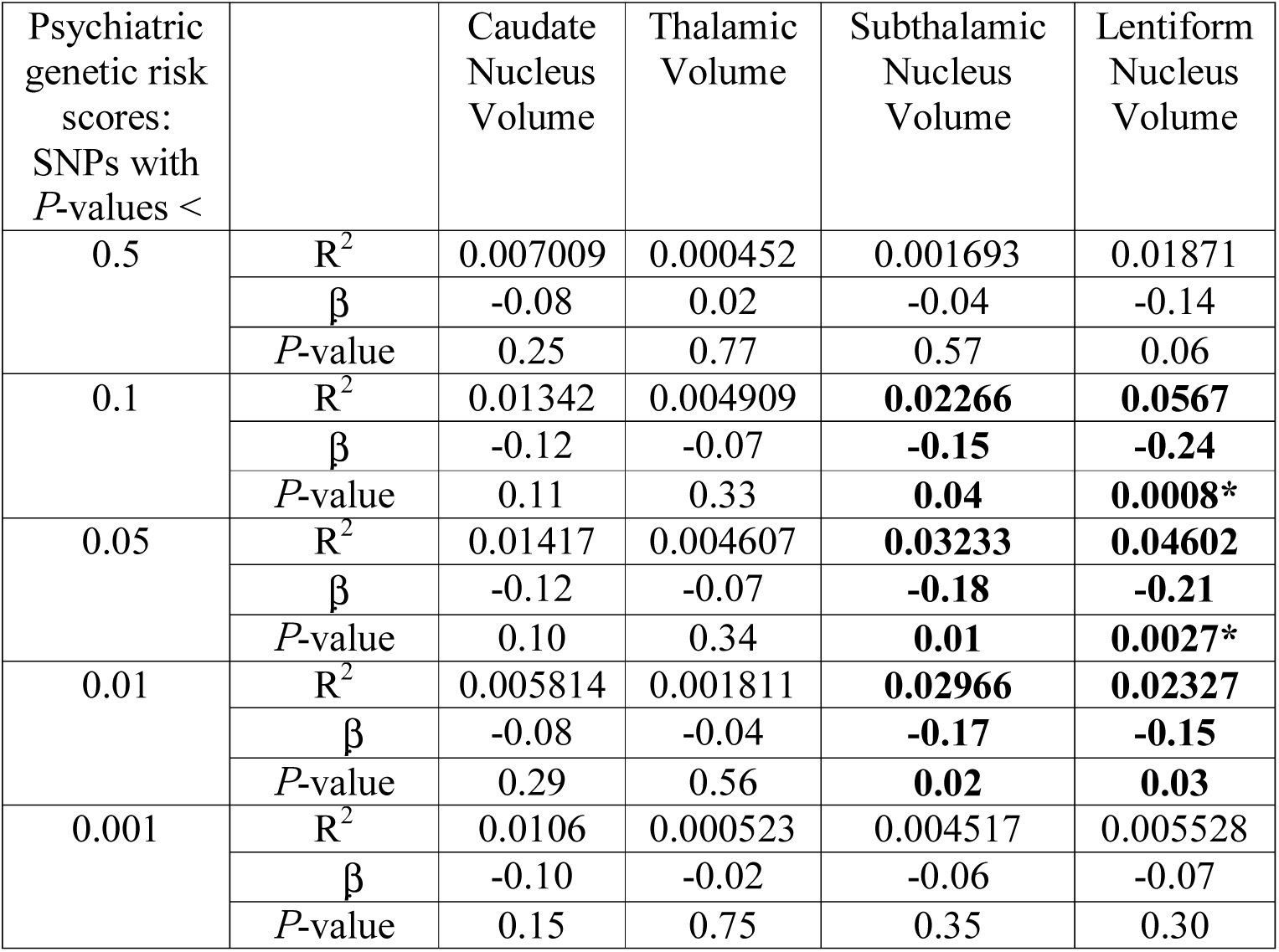
Effect size and significance of correlations between psychiatric PRS and deep grey matter volumes for the full mixed-ancestral cohort. Raw *P*-values are quoted. Raw *P*-values < 0.05 are shown in bold. Results surviving multiple-correction testing (p < 0.0083) are indicated with *.

**Table 3.** Effect size and significance of correlations between psychiatric PRS and deep grey matter volumes for the European cohort. Raw *P*-values are quoted. Raw *P*-values < 0.05 are shown in bold. Results surviving multiple-correction testing (p < 0.0083) are indicated with *.

| Psychiatric genetic risk scores: SNPs with $P$-values < | | Caudate Nucleus Volume | Thalamic Volume | Subthalamic Nucleus Volume | Lentiform Nucleus Volume |
| --- | --- | --- | --- | --- | --- |
| 0.5 | $R^2$ | 0.0114 | 8.451e-07 | 0.01263 | 0.02699 |
| | $\beta$ | -0.11 | -0.001 | -0.11 | -0.16 |
| | $P$-value | 0.24 | 0.99 | 0.22 | 0.07 |
| 0.1 | $R^2$ | 0.01247 | 0.01219 | **0.0669** | **0.05627** |
| | $\beta$ | -0.11 | -0.11 | **-0.26** | **-0.24** |
| | $P$-value | 0.22 | 0.23 | **0.004\*** | **0.008\*** |
| 0.05 | $R^2$ | 0.005469 | 0.003615 | **0.06896** | **0.04785** |
| | $\beta$ | -0.07 | -0.06 | **-0.26** | **-0.22** |
| | $P$-value | 0.42 | 0.51 | **0.003\*** | **0.015** |
| 0.01 | $R^2$ | 0.01572 | 0.004908 | **0.05949** | 0.02919 |
| | $\beta$ | -0.13 | -0.07 | **-0.24** | -0.17 |
| | $P$-value | 0.17 | 0.44 | **0.007\*** | 0.06 |
| 0.001 | $R^2$ | 0.005688 | 8.784e-05 | 0.007118 | 0.005267 |
| | $\beta$ | -0.07 | -0.01 | -0.08 | -0.07 |
| | $P$-value | 0.41 | 0.92 | 0.36 | 0.43 |

## 4. Results

Polygenic risk scores were computed at five different *P*-value thresholds (PT) for our sample of 194 preterm infants from summary statistics from the meta-analysis of genome-wide SNP data for five psychiatric disorders from the Cross-Disorder Group of the Psychiatric Genomics Consortium (20). These were compared with structural MRI brain measures of four deep grey matter volumes for our cohort of preterm infants. Sample characteristics are given in **Table 1**. These data were used to address the hypothesis that genetic variants which confer risk for psychiatric illness increase the chance of abnormal deep grey matter development following preterm birth.

The psychiatric PRS predicted subthalamic and lentiform nucleus volumes in preterm infants in both the full mixed-ancestral cohort and the sub-sample of European infants. In all cases, larger PRS were associated with smaller deep grey matter volumes.

For the full, mixed-ancestry cohort, the psychiatric PRS was negatively associated with lentiform nuclear volume (β = −0.24, p = 8×10-4, R^2^ = 0.057, (p_T_ = 0.1)); the psychiatric PRS also showed a modest negative association with subthalamic nuclear volume (β = −0.18, p = 0.01, R^2^ = 0.032, (p_T_ = 0.05)) which did not survive multiple testing correction (**Table 2 and Figures 1a. and 1b**). In the sub-sample of European infants, the psychiatric PRS was negatively associated with the lentiform nuclear volume (β = −0.24, p = 8×10^−3^, R^2^=0.056, (p_T_ =0.1)) and the subthalamic nuclear volume (β = −0.26, p = 3×10^−3^, R^2^=0.069, (p_T_ =0.05)) (**Table 3 and Figures 1c. and 1d**.). No association was found between the psychiatric PRS and caudate or thalamic volume for either the full mixed-ancestral sample or the sub-sample of European infants (**Table 2 and Table 3**). We note that the inclusion of postmenstrual age at scan as an additional regressor in the analysis did not significantly alter the results.

**Figure 1.**
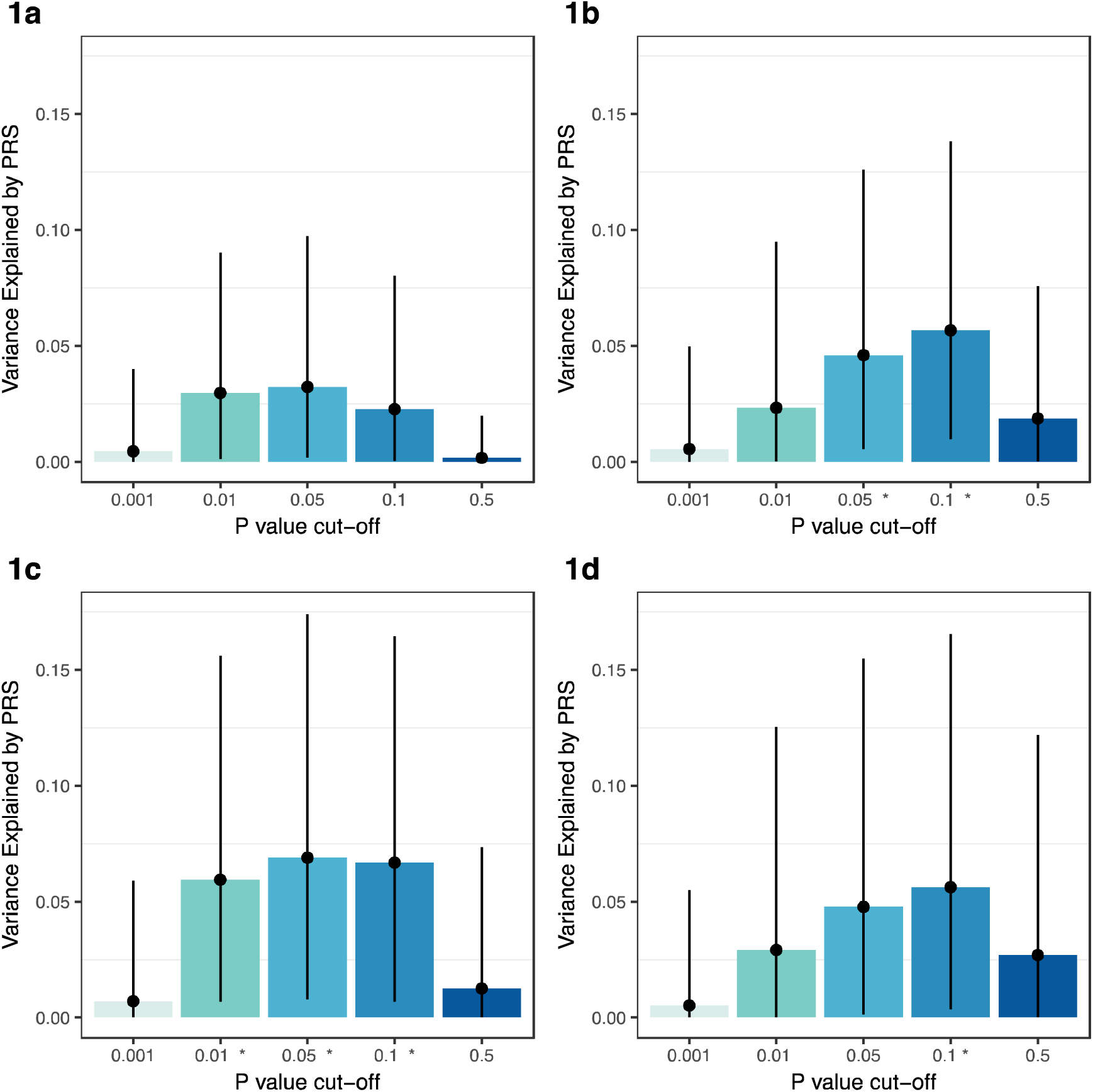
Proportion of variance explained in the subthalamic nucleus and lentiform nucleus volumes by the psychiatric PRS at five different *P*-value thresholds. Plots indicate the variance explained with estimated 95% confidence interval. The x-axis displays the five different upper thresholds of *P*-values for inclusion in the PRS. Results significant after multiple testing correction are indicated with an asterisk (*). **Figure 1a**. Subthalamic nucleus, full mixed-ancestral cohort, **Figure 1b**. Lentiform nucleus, full mixed-ancestral cohort, **Figure 1c**. Subthalamic nucleus, European sub-sample, **Figure 1d**. Lentiform nucleus, European sub-sample.

**Figure 2.**
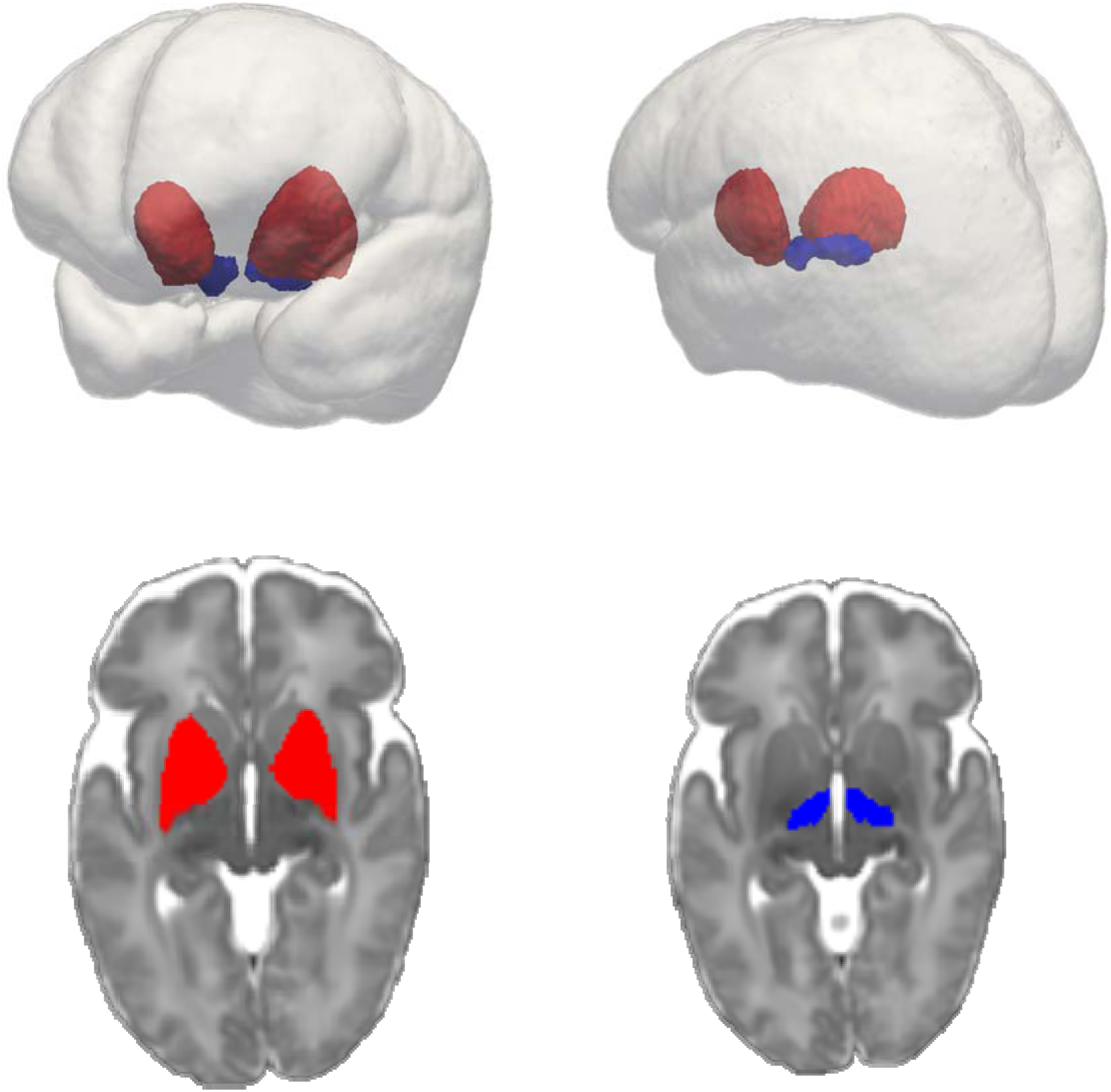
Subthalamic nucleus (blue) and Lentiform nucleus (red) within glass brain (top right and top left). Lentiform nucleus (bottom left) and subthalamic nucleus (bottom right) overlayed on 40-week neonatal template (axial cut).

As an additional check, we looked for a possible relationship between our psychiatric PRS and gestational age, no statistically significant relationship was found at any of the five *P*-value cut-offs. We also looked for a possible relationship between our psychiatric PRS and developmental outcome, assessed using scores from the five subtests within Bayley III: Cognitive, Expressive Communication, Receptive communication, Fine Motor and Gross Motor. A modest negative association was observed between Expressive Communication and psychiatric PRS (p = 0.013, β =−0.18, (P_T_=0.01)).

## 5. Discussion

We found evidence of an association between polygenic risk for psychiatric pathology and reduced lentiform volume in preterm infants. At its most predictive, in the full mixed-ancestry cohort, the PRS explained approximately 6% of the variance in the lentiform nucleus volume. This is a comparatively large effect for polygenic score analysis which typically explains only small amounts of variance (21).

This work employs summary statistics from the work of (20) using results from their overall analysis combining five psychiatric disorders: autism spectrum disorder (ASD), attention deficit-hyperactivity disorder (ADHD), bipolar disorder, major depressive disorder, and schizophrenia. Preterm infants have high levels of sub-diagnostic psychiatric symptomatologies and are at high risk of a broad range of psychiatric diagnoses, notably bipolar disorder, ADHD and ASD. The empirical evidence of shared genetic etiology for psychiatric disorders is strong. (22) showed a strong genetic correlation between schizophrenia and bipolar disorder (0.68 ± 0.04 s.e.), and moderate between schizophrenia and major depressive disorder (0.43 ± 0.06 s.e.), bipolar disorder and major depressive disorder (0.47 ± 0.06 s.e.), and ADHD and major depressive disorder (0.32 ± 0.07 s.e.). This commonalty in both symptomatology and genetic predisposition supports the use of a combined PRS in this study.

We hypothesised that common genetic risk variants which increase the risk of psychiatric pathology also increase vulnerability to aberrant development of deep grey matter structures in individuals subjected to the unusual environmental stresses associated with preterm birth.

It is possible that genetic variants associated with psychiatric risk predispose infants born preterm to abnormal lentiform development when such individuals are subjected to the unique environmental stresses of prematurity. An alternative explanation is that these variants are more ubiquitously associated with abnormal lentiform development and that this association is independent of the environmental precipitants associated with prematurity and would be similarly observed in a cohort of term-born infants or adults.

A number of studies have explored links between brain structure and genetic risk for psychiatric disorders in the adult population (23)(24)(25). (23) found polygenic risk for schizophrenia was significantly associated with total brain volume. Other authors (24) compared subcortical brain volume measures and PRS for schizophrenia and bipolar disorder in a sample of healthy adults; they found that most subcortical structures showed no association with a schizophrenia PRS with the exception of the globus pallidus and amygdala.

The most recent and largest study undertaken by (25) looked at the impact of polygenic risk of major depressive disorder, schizophrenia, and bipolar disorder on subcortical brain volume and white matter microstructure. They found no statistically significant associations between either subcortical volumes or white matter microstructure and psychiatric polygenic risk, although they note a modest negative association between thalamic volume and the polygenic risk for schizophrenia. (26) used a schizophrenia polygenic risk score and linkage disequilibrium (LD) score regression to test for shared genetic architecture between subcortical brain volume and schizophrenia and failed to find any overlap.

These large recent studies in adults have failed to show a robust association between deep grey matter and psychiatric risk (25)(26). This makes our second explanation, that genetic variants associated with psychiatric risk are more generally associated with abnormal lentiform development, independent of environmental pressures, less likely. However, associations between brain volume and psychiatric risk might be more difficult to detect in older cohorts, or in cohorts where the environmental stressor is less extreme than preterm birth. Brain volumes in older individuals have been subject to variable environmental exposures for a far greater time than neonatal brains. Environmental effects, which include the influence of psychotropic medication (27) may make genetic influences on brain structure more difficult to detect. It may be that the trajectory of growth of these structures is such that genetic overlap with psychiatric pathology is most easily detected early in life.

This study used an imaging endophenotype extracted from a large imaging dataset which was homogeneous in terms of MRI acquisition protocol, data pre-processing, analysis and quality control. The sample is small compared with many genetics studies, but endophenotypes have proved effective in allowing studies of small datasets, including in the perinatal period (28). Nevertheless, replication of these results will be required in independent samples.

Furthermore, our cohort is ancestrally diverse. The discovery GWAS sample used to derive the PRS is of European ancestry whereas our full cohort includes infants of European, African and Asian origin. Excluding non-European infants would have significantly reduced our sample size and it is also important to generate growing amounts of evidence across a variety of common ancestral populations. We therefore undertook our analysis both in the full mixed-ancestral sample and performed an additional sensitivity analysis in the sub-sample of European babies. The general stability of results across the two cohorts is reassuring.

Our hypothesis was motivated by studies in preterm infants showing abnormal deep grey matter development (6)(7) and its association with long-term neurodevelopmental outcome (9)(29). However, it is possible that other neuroimaging phenotypes both structural and functional could show a greater association with genetic risk for psychiatric pathology. Furthermore, there may be genetic variants associated with psychiatric disease influencing biology that is not easily captured by MRI.

In summary, our study reports an association between volume of the lentiform nucleus in preterm infants and genetic risk for psychiatric pathology. Further annotation of the shared genetic architecture and its associated biological pathways may shed light on potential therapeutic strategies.

### Data Availability

Data are available from the authors upon request, subject to approval of future uses by the National Research Ethics Service. The study was approved by the National Research Ethics Service, and written informed consent was given by all participating families.

## 6. Acknowledgements and Disclosures

Our thanks to the EPRIME collaborators who helped with the data collection for this study and to the children and families who participated in the study, and the nurses, doctors, and scientists who supported the project. This report uses data from independent research commissioned by the National Institute for Health Research (NIHR) and funded by the NIHR Programme Grants for Applied Research Programme (RP-PG-0707-10154.). The views and opinions expressed by authors in this publication are those of the authors and do not necessarily reflect those of the NHS, the NIHR, MRIC, CCF, NETSCC, the Programme Grants for Applied Research programme or the Department of Health. The work was also supported by the NIHR Biomedical Research Centers at Guy’s and St Thomas’ NHS Trust and the Wellcome Trust Centre for Medical Engineering at King’s College London. The authors report no biomedical financial interests or potential conflicts of interest.

## 7. **Author Contributions**

Author Contributions: Study conceptualization and design: H.C, S.S, G.Breen, R.P, A.D.E, data collection: S.C, J.H, Data analysis: H.C, S.S, G. Ball, A.V, A.S, M.K. Writing and Figure prep: H.C, S.S, A.D.E, Reviewing manuscript: H.C, M.K, S.S, G.Ball, A.V., A.S., S.C., J.H., G.Breen., R.P., A.D.E.

## 8. References

1. Blencowe H, et al. (2012) National, regional, and worldwide estimates of preterm birth rates in the year 2010 with time trends since 1990 for selected countries: A systematic analysis and implications. Lancet 379(9832):2162–2172.

2. Moore T, et al. (2012) Neurological and developmental outcome in extremely preterm children born in England in 1995 and 2006: the EPICure studies. Bmj 345:e7961.

3. Mackay DF, Smith GCS, Dobbie R, Pell JP (2010) Gestational age at delivery and special educational need: Retrospective cohort study of 407,503 schoolchildren. PLoS Med 7(6):1–10.

4. Johnson S, Marlow N (2011) Preterm birth and childhood psychiatric disorders. Pediatr Res 69(5 PART 2):22–28.

5. Nosarti C, et al. (2012) Preterm birth and psychiatric disorders in young adult life. ArchGenPsychiatry 69(1538–3636 (Electronic)):E1--E8.

6. Boardman JP, et al. (2006) Abnormal deep grey matter development following preterm birth detected using deformation-based morphometry. Neuroimage 32(1):70–78.

7. Srinivasan L, Dutta R, Counsell SJ, Allsop JM, Boardman JP (2007) Quantification of Deep Gray Matter in Preterm Infants at Term-Equivalent Age Using Manual Volumetry of 3-Tesla Magnetic Resonance Images. 119(4). doi:10.1542/peds.2006-2508.

8. Ball G, et al. (2012) The effect of preterm birth on thalamic and cortical development. Cereb Cortex 22(5):1016–1024.

9. Inder TE, Warfield SK, Wang H, Hu PS (2005) Abnormal Cerebral Structure Is Present at Term in Premature Infants. 115(2). doi: 10.1542/peds.2004-0326.

10. Ball G (2017) Multimodal image analysis of clinical influences on preterm brain development⁪: Clinical factors and preterm brain … Multimodal image analysis of clinical influences on preterm brain development. (July). doi: 10.1002/ana.24995.

11. Dempfle A, et al. (2008) Gene-environment interactions for complex traits: definitions, methodological requirements and challenges. Eur J Hum Genet 16(10):1164–72.

12. Leviton A, Gressens P, Wolkenhauer O, Dammann O (2015) Systems approach to the study of brain damage in the very preterm newborn. Front Syst Neurosci 9(April):58.

13. Polderman TJC, et al. (2015) Meta-analysis of the heritability of human traits based on fifty years of twin studies. Nat Genet 47(7):702–709.

14. Kochunov P, et al. (2015) Heritability of fractional anisotropy in human white matter: a comparison of Human Connectome Project and ENIGMA-DTI data. Neuroimage 111:300–311.

15. Lin GG, Scott JG (2012) NIH Public Access. 100(2):130–134.

16. Gilmore JH, et al. (2010) Genetic and environmental contributions to neonatal brain structure: A twin study. Hum Brain Mapp 31(8):1174–1182.

17. Gratten J, Wray NR, Keller MC, Visscher PM (2014) Large-scale genomics unveils the genetic architecture of psychiatric disorders. Nat Neurosci 17(6):782–790.

18. Manolio TA, et al. (2009) Finding the missing heritability of complex diseases. Nature 461(7265):747–53.

19. Dudbridge F (2013) Power and Predictive Accuracy of Polygenic Risk Scores. PLoS Genet 9(3). doi:10.1371/journal.pgen.1003348.

20. Smoller JW (2013) Identification of risk loci with shared effects on five major psychiatric disorders: A genome-wide analysis. Lancet 381(9875):1371–1379.

21. Lee SH, et al. (2012) NIH Public Access. Nat Genet 44(3):247–250.

22. Lee SH, et al. (2013) Genetic relationship between five psychiatric disorders estimated from genome-wide SNPs. Nat Genet 45(9):984–994.

23. Terwisscha Van Scheltinga AF, et al. (2013) Genetic schizophrenia risk variants jointly modulate total brain and white matter volume. Biol Psychiatry 73(6):525–531.

24. Caseras X, Tansey KE, Foley S, Linden D (2015) Association between genetic risk scoring for schizophrenia and bipolar disorder with regional subcortical volumes. Transl Psychiatry 5(12):e692.

25. Reus LM, et al. (2017) Association of polygenic risk for major psychiatric illness with subcortical volumes and white matter integrity in UK Biobank. Nat Publ Gr (February): 1–8.

26. Franke B, et al. (2016) Genetic influences on schizophrenia and subcortical brain volumes: large-scale proof of concept. Nat Neurosci 19(3):420–431.

27. Moncrieff J, Leo J (2010) A systematic review of the effects of antipsychotic drugs on brain volume. Psychol Med 40(9):1409–1422.

28. Krishnan ML, et al. (2016) Possible relationship between common genetic variation and white matter development in a pilot study of preterm infants. Brain Behav 434:1–14.

29. Boardman JP, et al. (2010) A common neonatal image phenotype predicts adverse neurodevelopmental outcome in children born preterm. Neuroimage 52(2):409–414.

30. Tustison NJ, et al. (2010) N4ITK: Improved N3 Bias Correction. IEEE Trans Med Imaging 29(6):1310–1320.

31. Serag A, et al. (2012) Construction of a consistent high-definition spatio-temporal atlas of the developing brain using adaptive kernel regression. Neuroimage 59(3):2255–2265.

32. Rueckert D, Frangi AF, Schnabel JA (2003) Automatic construction of 3-D statistical deformation models of the brain using nonrigid registration. Med Imaging, IEEE Trans 22(8):1014–1025.

33. Makropoulos A, et al. (2016) Regional growth and atlasing of the developing human brain. Neuroimage 125:456–478.

34. Purcell S, et al. (2007) PLINK: A Tool Set for Whole-Genome Association and Population-Based Linkage Analyses. Am J Hum Genet 81(3):559–575.

35. Chang CC, et al. (2015) Second-generation PLINK: rising to the challenge of larger and richer datasets. Gigascience 4(1):7.

36. Euesden J, Lewis CM, O’Reilly PF (2014) PRSice: Polygenic Risk Score software. Bioinformatics 31(9):1466–1468.

37. Bayley N (2006) Bayley scales of infant and toddler development.

38. Li J, Ji L (2005) Adjusting multiple testing in multilocus analyses using the eigenvalues of a correlation matrix. Heredity (Edinb) 95(3):221–227.

